# Cryptochromes-mediated Inhibition of the CRL4^Cop1-^ Complex Assembly Defines an Evolutionary Conserved Signaling Mechanism

**DOI:** 10.1101/545533

**Authors:** Luca Rizzini, Daniel C. Levine, Mark Perelis, Joseph Bass, Clara B. Peek, Michele Pagano

## Abstract

In plants, Cryptochromes are photoreceptors that negatively regulate CRL4^Cop1^(Cul4-RING ubiquitin ligase and its substrate receptor Cop1). In mammals, Cryptochromes are core components of the circadian clock and repressors of the glucocorticoid receptor (GR). Moreover, mammalian Cryptochromes lost their ability to interact with Cop1, suggesting that they are unable to inhibit CRL4^Cop1^. Contrary to this assumption, we found that mammalian Cryptochromes are also negative regulators of CRL4^Cop1^and through this mechanism they repress the GR transcriptional network both in cultured cells and in mouse liver. Mechanistically, Cryptochromes inactivate Cop1 by interacting with Det1, a specific subunit of the mammalian CRL4^Cop1^complex. Through this interaction, the ability of Cop1 to join the CRL4 complex is inhibited; therefore, its substrates accumulate. Thus, the interaction between Cryptochromes and Det1 in mammals mirrors the interaction between Cryptochromes and Cop1 *in planta,* pointing to a common ancestor in which the cryptochrome-Cop1 axis originated.

## Main Text

Plant Cryptochromes are UVA- and blue-light photoreceptors. One of their major functions, after UVA- and blue-light perception, is to inactivate the ubiquitin ligase activity of Cop1 (Constitutively photomorphogenic 1) [1, 2]. Cop1 is a DCAF (Ddb1 and Cul4-Associated Factor) protein that functions as a substrate receptor subunit of the CRL4^Cop1^ complex, which is comprised of the following five subunits: Cul4, Ddb1, Rbx1, Dda1, and Cop1 [3–5]. In plants, CRL4^Cop1^promotes the ubiquitylation and the consequent proteasomal-dependent degradation of downstream substrates, such as the photomorphogenesis-promoting transcription factors HY5, HYH and LAF1 [1]. Cryptochrome-mediated inactivation of the CRL4^Cop1^complex results in the stabilization of these factors and contribute to the reprogramming of plant growth and development [1].

Mammalian Cryptochromes, Cry1 and Cry2, are instead core repressors of the circadian clock [6, 7] and, in contrast to their plant orthologs, do not interact with Cop1 [8]. Cryptochromes, the Period proteins, and Rev-Erb ⍰, constitute the negative limb of the transcription-translation feedback loop (TTFL) by binding and inhibiting the Clock/Bmal1 complex within the forward limb of the clock. In addition, mammalian Cryptochromes regulate the activity of the glucocorticoid receptor (GR) and modulate cAMP signaling pathways important in gluconeogenesis [9, 10].

Cop1 is also present in mammals where it functions as a DCAF protein allowing the CRL4^Cop1^complex to recognize specific substrates [1]. In particular, mammalian Cop1 promotes the ubiquitylation and subsequent proteasomal-dependent degradation of downstream substrates (mostly transcription factors and metabolic enzymes), such as c-Jun, p53, Mta1, Stat3, Ets1, and Atgl [3, 11–15]. Thus, Cop1 constitutes a hub that rapidly modulates the stability of key cellular regulators in response to flux of nutrients and hormones, as well as upon cellular stress.

### Cop1 mediates the Cryptochromes-dependent rhythmic repression of the GR

Given the lack of physical interaction between mammalian Cryptochromes and Cop1 [8], it is assumed that the Cryptochromes-Cop1 axis is not evolutionary conserved. However, functional studies to investigate the presence of this regulatory axis in mammals have not been performed. As a first step towards this goal, we investigated the involvement of Cop1 in the well-established function of the mammalian Cryptochromes in repressing the GR transcriptional network [9]. Compared to wild-type mouse embryonic fibroblasts (MEFs), Cry1^-/-^;Cry2^-/-^ double knockout (DKO) MEFs display an elevated level of GR activity [9]. We expanded these findings by exogenously expressing either Cry1 or Cry2 using a doxycycline-inducible system (TET-ON), in DKO MEFs. After activation of the GR pathway with dexamethasone (a synthetic glucocorticoid), we noticed that the expression of *Sgk1* and *Gilz,* two canonical GR target genes, was reduced upon induction of either Cry1 or Cry2 (Fig. S1A-B).

To understand whether a genetic interaction between Cryptochromes and Cop1 is present in mammals, we measured dexamethasone-dependent activation of *Sgk1* and *Gilz* upon depletion of Cop1. We found that Cop1 silencing inhibited GR-induced transcription of *Sgk1* and *Gilz* (Fig. 1A and Fig. S1C). Moreover, in cells depleted of Cop1, induction of Cry1 or Cry2 had no additional effect on the repression of *Sgk1* and *Gilz* (Fig. 1A and Fig. S1C), suggesting that Cryptochromes and Cop1 operate in the same GR-repressing signaling pathway.

**Fig. 1.**
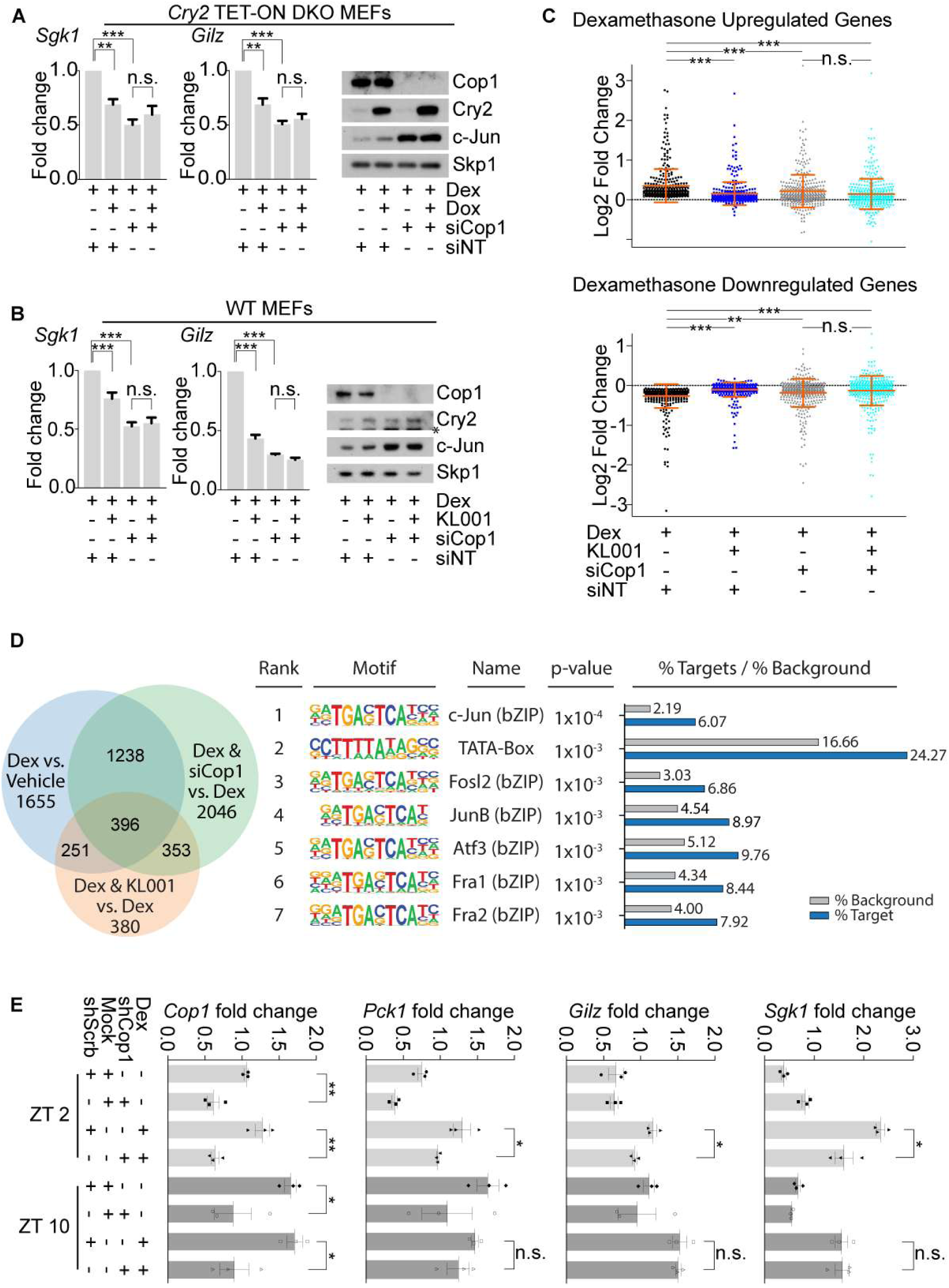
Cop1 mediates the Cryptochromes-dependent rhythmic repression of the GR. **(A)** On the left, qPCR analysis of cDNA prepared from DKO MEFs infected with lentiviruses expressing untagged Cry2 under the control of a doxycycline-inducible promoter, after dexamethasone treatment, in presence or absence of Cop1 knockdown. On the right, cells extracts were analyzed by immunoblotting with antibodies to the indicated proteins. Human Cry2 was exogenously expressed in DKO MEFs and tested with antibodies against human Cry2. The Skp1 protein has been used as a loading control. **(B)** On the left, qPCR analysis of cDNA prepared from wild-type MEFs, before and after KL001 treatment, after dexamethasone treatment and in presence or absence of Cop1 knockdown. On the right, cells extracts were analyzed by immunoblotting with antibodies to the indicated proteins. The asterisk indicates a cross-reacting band. **(C)** Dexamethasone-mediated activation of the GR in wild-type MEFs treated with KL001, in presence or absence of Cop1 knockdown. RNA was isolated from the samples and quantified by RNA-seq. Differential expression analysis of RNA-seq data sets, [see next-generation sequencing data deposited in Gene Expression Omnibus (GEO) database under the accession code GSE124388] is represented in the scatter plots as log2 fold change for 792 genes divided in Up- and Down-regulated. Dexamethasone-regulated genes were considered differentially expressed when the p-adjusted value was <0.05. Hence, 370 Up-regulated & 422 Down-regulated genes were displayed. (n=3 biologically independent experiments). **(D)** MEFs were treated with dexamethasone either alone or in combination with KL001 treatment or Cop1 depletion. In the left panel, Venn diagram represents all the genes significantly differentially expressed among al tested conditions. RNA was isolated from the samples and quantified by RNA-seq. [see next-generation sequencing data deposited in Gene Expression Omnibus (GEO) database under the accession code GSE124388]. Regulated genes were considered differentially expressed when the p-adjusted value was <0.05. In the right panel, the table contains the first 7 most highly overrepresented motif within the promoter regions of the 396 commonly regulated genes among all the tested conditions. **(E)** Mice were retro-orbitally injected with Adeno-associated viruses expressing either scramble shRNA or Cop1 shRNA. After 4 weeks, mice were tail-injected with dexamethasone for 1 hour and then sacrificed at ZT2 and ZT10. Liver tissue was then collected, and RNA was extracted and analyzed by qPCR.

To analyze the function of the Cryptochrome-Cop1 axis in GR signaling in wild-type MEFs, we stabilized endogenous Cryptochromes using the carbazole derivative KL001. KL001 competes with the CRL1^Fbxl3^, a Cul1-RING ubiquitin ligase complex, for its binding to Cryptochromes, thereby resulting in the inhibition of their ubiquitylation and degradation [16–19]. First, we noticed that KL001 repressed the dexamethasone-mediated induction of *Sgk1* and *Gilz*, in agreement with the accumulation of Cry2 in KL001-treated cells (Fig. S1D). Moreover, in cells depleted of Cop1, KL001 treatment had no additive effect on the expression of *Sgk1* and *Gilz* (Fig. 1B). These data are in agreement with the results obtained with the induction of exogenous Cryptochromes in DKO MEFs and, together, they suggest that, in the absence of Cop1, Cryptochromes are impaired in their ability to inhibit GR. Moreover, these results indicate that KL001 acts through a novel pathway involving Cop1 to inhibit GR activity.

Next, we performed differential expression analysis using RNA-sequencing to determine the impact of the genetic interactions between Cryptochromes, Cop1, and GR on a genomic scale. Dexamethasone-mediated activation of the GR was analyzed in wild-type MEFs treated with KL001 (to stabilize the Cryptochromes) in the presence or in the absence of Cop1 knockdown. We found that out of 3,540 genes significantly differentially expressed by dexamethasone, 792 lost significant expression after KL001 treatment [see next-generation sequencing data deposited in Gene Expression Omnibus (GEO) database under the accession code GSE124388]. The scatter plots and the heatmap representations of these 792 genes shows that KL001 treatment blocked the effects of dexamethasone on both up- and down-regulated genes (Fig. 1C and Fig. S1E). Importantly, in cells depleted of Cop1, KL001 treatment had no significant effect on these 792 GR-regulated genes (Fig. 1C and Fig. S1E). Altogether, these results indicate that Cop1 is a regulator of GR signaling and a downstream effector of the Cryptochromes-mediated repression of the GR transcriptional activity.

We also noticed that 396 GR-regulated genes were significantly differentially expressed when dexamethasone was provided either together with KL001 or in Cop1 knockdown cells (Fig. 1D, left panel). Motif enrichment analysis for these 396 genes showed that the first 7 most highly overrepresented motifs within the promoter regions of these genes contained the core sequence TGAC/GTCA which is the recognition site for AP-1 transcription factors (Fig. 1D, right panel). The AP-1 specific signature was absent from the remaining dexamethasone-dependent gene promoters identified in this analysis (Fig. S1F) suggesting that part of the transcriptional output downstream of Cop1 and KL001 is regulated by c-Jun. This is noteworthy since c-Jun is both a constitutive substrate of Cop1 and (in complex with Fos family members, such as Fra1 and Fra2) a repressor of the GR transcriptional activity [20–23].

Cryptochromes mediate the rhythmic repression of GR in mouse liver, displaying its maximal and minimal response to dexamethasone at ZT2 and ZT10, respectively [9]. We confirmed and expanded these findings by showing that the expression of the GR target genes *Sgk1*, *Gilz,* and *Pck1* is inhibited in mouse liver specifically at ZT10, but not at ZT2 (Fig. 1E). To test whether time-of-day dependent GR target gene activation is dependent on Cop1, we depleted Cop1 *in vivo* using an adenoviral-mediated shRNA injected retro-orbitally in mice. Confirming our data in cell systems, *in vivo* depletion of Cop1 was found to inhibit the dexamethasone-mediated activation of GR (Fig. 1E). Cop1 depletion inhibited specifically the GR activation at ZT2 (Fig. 1E), when the GR is not repressed by the Cryptochromes [9]. Instead, Cop1 depletion at ZT10 did not additionally inhibit the activation of the GR, at a time when GR transcriptional activity is already repressed by the Cryptochromes. Again, these data confirm our results in MEFs where the Cryptochromes- and Cop1-mediated transcriptional repression of GR are not additive (Fig. 1A-C, Fig. S1C, and Fig. S1E).

### Cry2 controls the level and stability of Cop1’s substrates

The above results revealed the existence of a genetic interaction among Cryptochromes and Cop1 that encouraged us to investigate the underlying mechanism. We noticed that the protein levels of c-Jun and JunB, two canonical Cop1 substrates, were lower in *Cry1/2* DKO MEFs than those present in MEFs isolated from wild-type littermate mice (Fig. 2A).

**Fig. 2.**
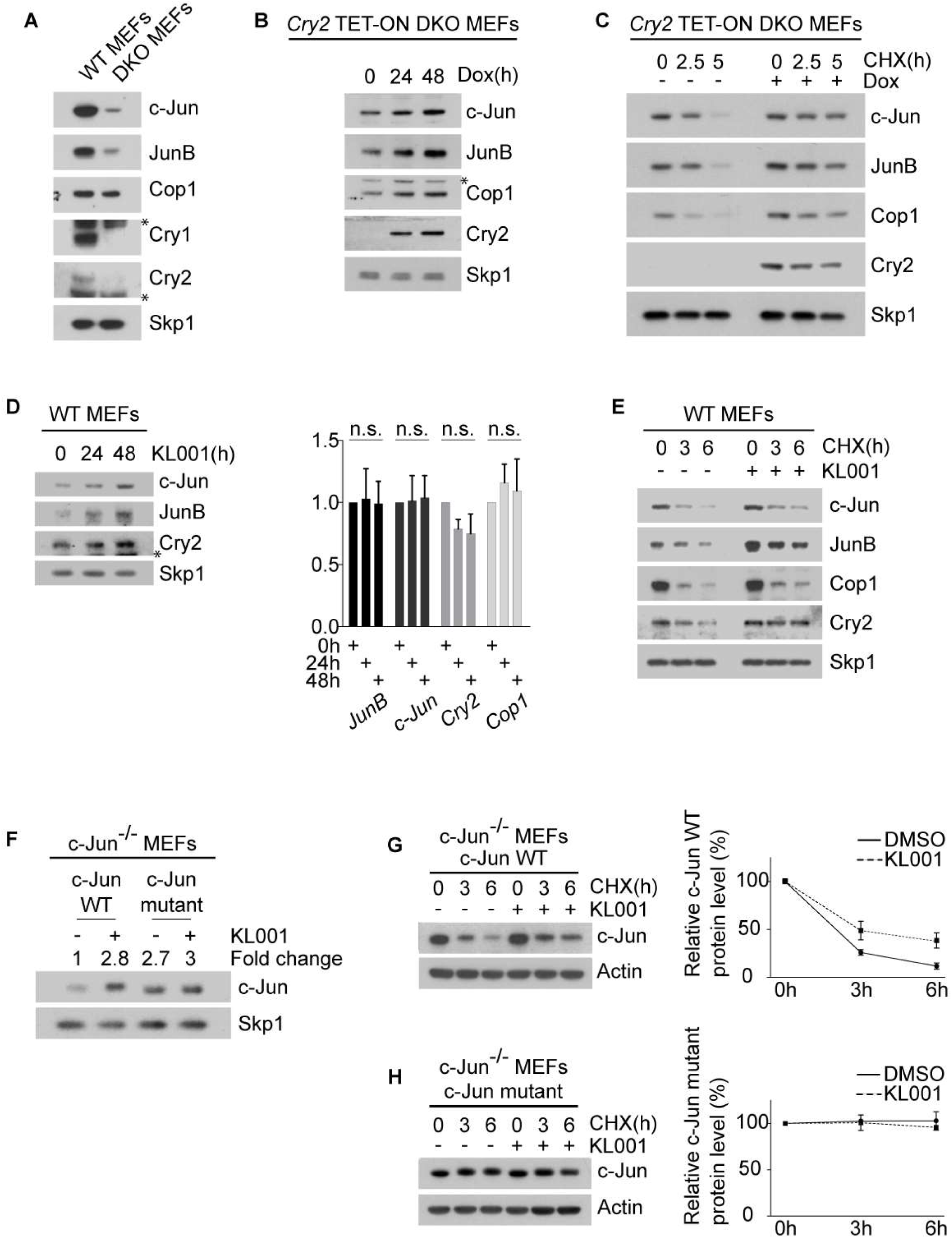
Cry2 controls the level and stability of Cop1’s substrates. **(A)** Cell extracts of wild-type MEFs and DKO MEFs were analyzed by immunoblotting with antibodies to the indicated proteins. The asterisks indicate cross-reacting bands. **(B)** Western blot analysis of Cop1’s substrates in DKO MEFs infected with lentiviruses expressing untagged Cry2 under the control of a doxycycline-inducible promoter. After doxycycline treatment, cells were collected, lysed and immunoblotted with antibodies to the indicated proteins. Human Cry2 was exogenously expressed in DKO MEFs and tested with antibodies against human Cry2. The asterisk indicates a cross-reacting band. **(C)** DKO MEFs infected with lentiviruses expressing untagged Cry2 under the control of a doxycycline-inducible promoter were treated overnight with doxycycline. The next day cycloheximide was applied to the cells for the indicated time and cells were then collected, lysed and immunoblotted with antibodies to the indicated proteins. Human Cry2 was exogenously expressed in DKO MEFs and tested with antibodies against human Cry2. **(D)** On the left, wild-type MEFs were treated with KL001 10 [µM] for the indicated time before sample collection. Cell extracts were analyzed by immunoblotting with antibodies to the indicated proteins. The asterisk indicates a cross-reacting band. On the right, qPCR analysis of cDNA prepared from the same samples. **(E)** Wild-type MEFs were treated with KL001 10 [µM] for 18 hours. Cells were then treated with cycloheximide for the indicated time and cells extracts were analyzed by immunoblotting with antibodies to the indicated proteins. **(F)** c-Jun^-/-^ MEFs stably expressing either c-Jun wild-type or a degron mutant of c-Jun were treated for 2 days with KL001 10 [µM]. Cells extracts were then analyzed by immunoblotting with antibodies to the indicated proteins. The fold change indicates the protein levels of c-Jun wild-type and c-Jun mutant after densitometric analysis of the blot. **(G)** c-Jun^-/-^ MEFs stably expressing c-Jun wild-type were treated overnight with KL001 10 [µM] where indicated. Cycloheximide was applied for the indicated time and cells extracts were analyzed by immunoblotting with antibodies to the indicated proteins. On the right, the quantification of c-Jun wild-type protein levels ±SEM (n = 3 biologically independent experiments). **(H)** c-Jun^-/-^ MEFs stably expressing a c-Jun mutant in its Cop1’s degron were treated overnight with KL001 10 [µM] where indicated. Cycloheximide was applied for the indicated time and cells extracts were analyzed by immunoblotting with antibodies to the indicated proteins. On the right, the quantification of c-Jun degron mutant protein levels ±SEM (n = 3 biologically independent experiments).

Similarly, Cop1, which is known to induce its own degradation [24], displayed lower levels in DKO MEFs compared to wild-type MEFs. Notably, doxycycline-mediated expression of Cry1 or Cry2 in DKO MEFs increased the levels of Cop1, c-Jun, and JunB over time (Fig. 2B and Fig. S1A-B). These findings mirror what was observed in Cryptochromes knockout *Arabidopsis* seedlings, where the transcription factor HY5, a canonical substrate of Cop1 in plants, displays lower protein levels than in wild-type seedlings [25], but increases upon expression of exogenous Cryptochromes [26].

We also observed that the half-life of Cop1, c-Jun, and JunB increased after induction of exogenous Cry2, showing that Cry2 modulates their protein stability (Fig. 2C). Similar protein stabilization was observed for various Cop1 substrates in all cell systems tested, including mouse Beta-TC-6 pancreatic beta cells and human HEK-293T embryonic kidney cells (Fig. S2). Accordingly, KL001, which stabilizes Cryptochromes, induced the stabilization of Cop1 substrates without affecting their corresponding mRNAs (Fig. 2D-E, Fig. 1B, and Fig. S1D).

To uncouple post-translational regulation of protein stability from endogenous transcriptional regulation and other RNA regulatory elements, we also analyzed in c-Jun^-/-^ MEFs the protein level and the stability of exogenously-expressed c-Jun [either wild-type c-Jun or a c-Jun stable mutant unable to bind Cop1 [3]]. We found that KL001 increased both the levels and stability of exogenous wild-type c-Jun (Fig. 2F-G). Instead, the c-Jun Cop1-insensitive stable mutant was not further stabilized by KL001 treatment (Fig. 2H). As expected, induction of either Cry1 or Cry2, as well as treatment with KL001 had no additive effect on the levels of c-Jun in Cop1-depleted MEFs (Fig. 1A-B and Fig. S1C). Thus, Cryptochromes appear to promote the stabilization of c-Jun in a Cop1-mediated fashion.

### The N-terminus of Cry2 interacts with Det1 to stabilize Cop1’s substrates and to inhibit GR

The association between Cop1 and Cul4 is essential to form an active CRL4^Cop1^ubiquitin ligase complex and to mediate the ubiquitylation of downstream substrates [3]. Other subunits of the CRL4^Cop1^complex include Ddb1, which is also present in all CRL4 complexes, and Dda1, which is present in most but not all CRL4 complexes [4]. Moreover, CRL4^Cop1^is the only mammalian CRL4 complex known to contain an additional subunit, namely Det1, which plays an essential role as an assembly factor to link Cop1 to Ddb1 [3]. Using a panel of eight DCAF proteins, we confirmed that Det1 specifically binds Cop1 (Fig. S3A). We also confirmed that Cop1 associates more strongly with Cul4A than with its paralog Cul4B, as previously observed [27]. Importantly, when we expressed increasing amounts of Det1 in HEK-293T cells, we observed that Cop1 co-immunoprecipitated increasing amounts of Cul4A, Dda1, and Ddb1 (Fig. S3B), suggesting that Det1 is a rate-limiting factor for Cop1 to associate with the rest of the CRL4 complex.

While in mammals Det1 is essential for Cop1 binding to Ddb1 and to form an active CRL4^Cop1^complex, plant Det1 is not present in the CRL4^Cop1^complex formation and it assembles instead in a distinct ligase complex, namely CRL4^Det1^ [1, 3, 5]. Moreover, plant Cryptochromes physically interact and inhibit Cop1, whereas in mammals this physical interaction is lost [8]. Because of these differences, we tested the hypothesis that mammalian Cryptochromes, instead of binding Cop1, could bind Det1.

Immunoprecipitation of HA-tagged Det1 expressed in HEK-293T cells revealed an interaction between Det1 and wild-type Cry2, which was tagged at the C-terminus with a FLAG epitope (Fig. 3A). We additionally noticed that adding a FLAG tag to the N-terminus of the two Cryptochrome proteins impaired their binding to Det1 (data not shown), suggesting that the N-terminal portion of the Cryptochrome is important for the binding to Det1. Therefore, we generated C-terminal deletion mutants of Cry2, all C-terminally tagged with a FLAG epitope (see schematics in Fig. 3B) and tested them for their binding to Det1. Immunopurification of HA-tagged Det1 co-expressed with either wild-type Cry2 or Cry2 truncation mutants revealed that the first N-terminal 130 amino acids of human Cry2 were sufficient for binding to Det1 (Fig. 3A-B). This is also divergent from the mechanism *in planta* where Cryptochromes bind and inhibit Cop1 through their C-terminal domain [26, 28]. Instead, the C-terminus of the mammalian Cryptochromes is critical for the interaction with components of the circadian clock, such as the Period proteins and Fbxl3 [29–31]. Hence, it was not surprising that the Cry2-N130 truncation mutant, although able to interact with endogenous Det1, Dda1, and Ddb1, lost its ability to bind with the components of the circadian clock (Fig. 3C).

**Fig. 3.**
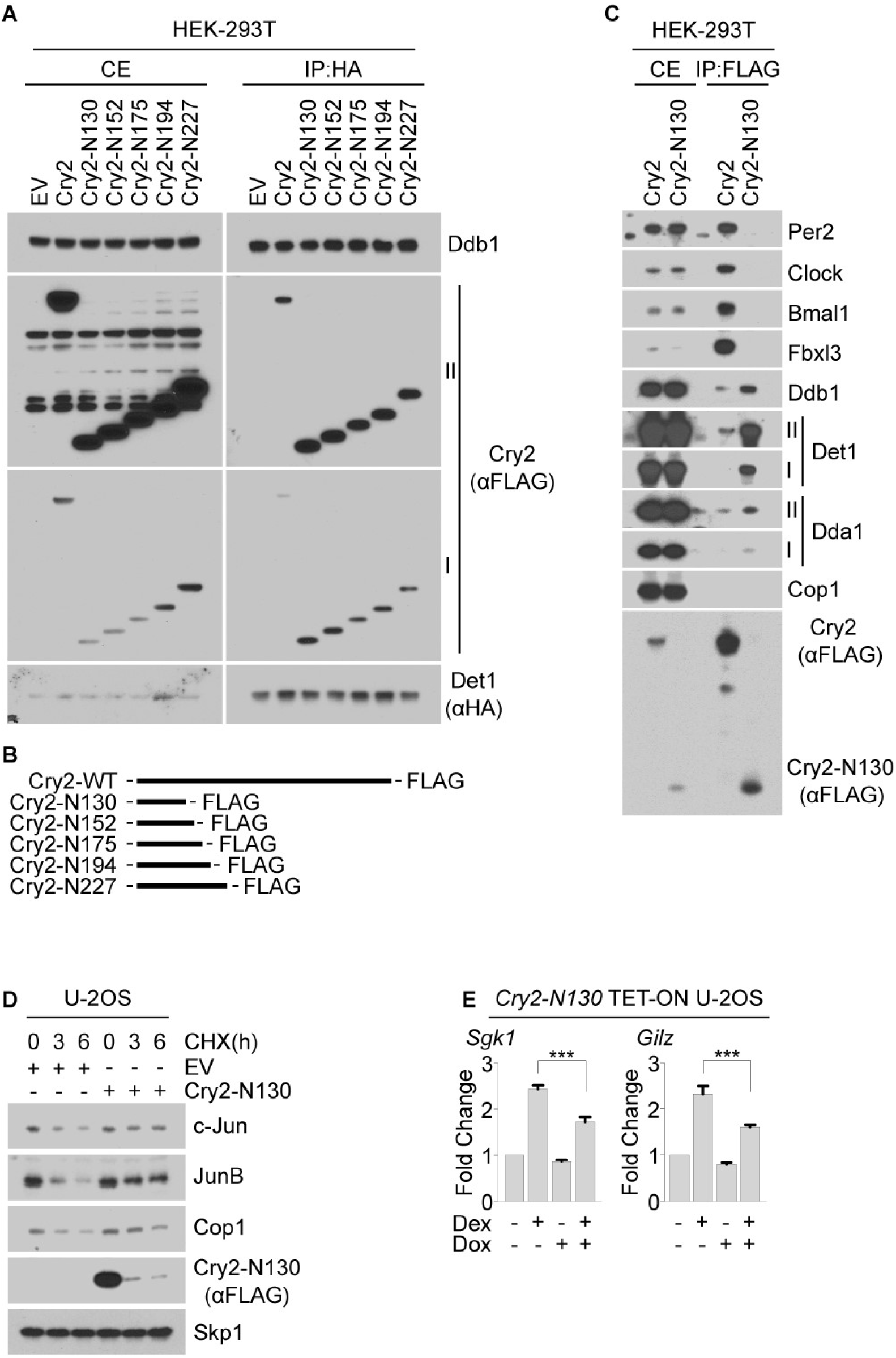
The N-terminus of Cry2 interacts with Det1 to stabilize Cop1’s substrates and inhibit GR activity. **(A)** Exogenously expressed HA tagged Det1 co-expressed with either FLAG tagged Cry2 wild-type or FLAG tagged Cry2 truncation mutants in HEK-293T cells followed by immunoprecipitation from cell extracts with anti-HA resin. Cells extracts and immunoprecipitates were analyzed by immunoblotting with antibodies to the indicated proteins. Roman numbers indicate different exposure times: I, short exposure; II, long exposure. **(B)** Schematic representation of Cry2 wild-type and Cry2 truncation mutants. **(C)** Exogenously expressed FLAG tagged Cry2 wild-type or FLAG tagged Cry2-N130 truncation mutant were expressed in HEK-293T cells followed by immunoprecipitation from cell extracts with anti-FLAG resin. Cell extracts and immunoprecipitates were analyzed by immunoblotting with antibodies to the indicated proteins. Roman numbers indicate different exposure times: I, short exposure; II, long exposure. **(D)** U-2OS cells were transfected with either GFP expressing vector or with a vector expressing the Cry2 truncation mutant, Cry2-N130. 18 hours after transfection, the cells were treated with CHX for the indicated time and cell extracts were collected and immunoblotted with antibodies to the indicated proteins. **(E)** qPCR analysis of cDNA prepared from U-2OS cells infected with lentiviruses expressing the Cry2 truncation mutant Cry2-N130 under the control of a doxycycline-inducible promoter and treated with dexamethasone where indicated.

Notably, expression of the truncation mutant Cry2-N130 in U-2OS cells was sufficient to increase the half-life of Cop1 substrates (Fig. 3D) and to reduce the dexamethasone-dependent activation of *Sgk1* and *Gilz* (Fig. 3E), similar to what we observed with full length Cryptochromes (Fig. 1 and Fig. 2). These data uncover a novel function of the N-terminal region of the mammalian Cryptochromes that is independent from their interactions with other components of the TTFL.

### Cry1 and Cry2 inhibit the binding between Cop1 and Det1-Ddb1

Next, to gain insights into the role of Cryptochromes in the assembly of the CRL4^Cop1^ complex, we immunopurified FLAG-tagged Cop1 expressed in HEK-293T cells either alone or in combination with Cry1 or Cry2. Mass-spectrometric analysis of the proteins present in the Cop1 complex showed that, when either Cry1 or Cry2 were co-expressed with Cop1, there was a reduction in the number of peptides corresponding to endogenous Dda1, Ddb1, and Det1 (Fig. S4A). To confirm these data, we tested the interaction between Cop1 and Ddb1 in both wild-type and DKO MEFs. Strikingly, immunopurification of endogenous Cop1 revealed a higher level of binding to endogenous Det1 and Ddb1 in DKO MEFs compared to wild-type MEFs (Fig. 4A). Additionally, expression of exogenous Cry2 in DKO MEFs reduced the binding of endogenous Cop1 to endogenous Det1 and Ddb1 (Fig. 4B).Similarly, transient expression of either Cry1 or Cry2 reduced the binding of exogenous Cop1 to endogenous Det1 and Ddb1 without affecting the interaction between COP1 and c-Jun (Fig. S4B).

**Fig. 4.**
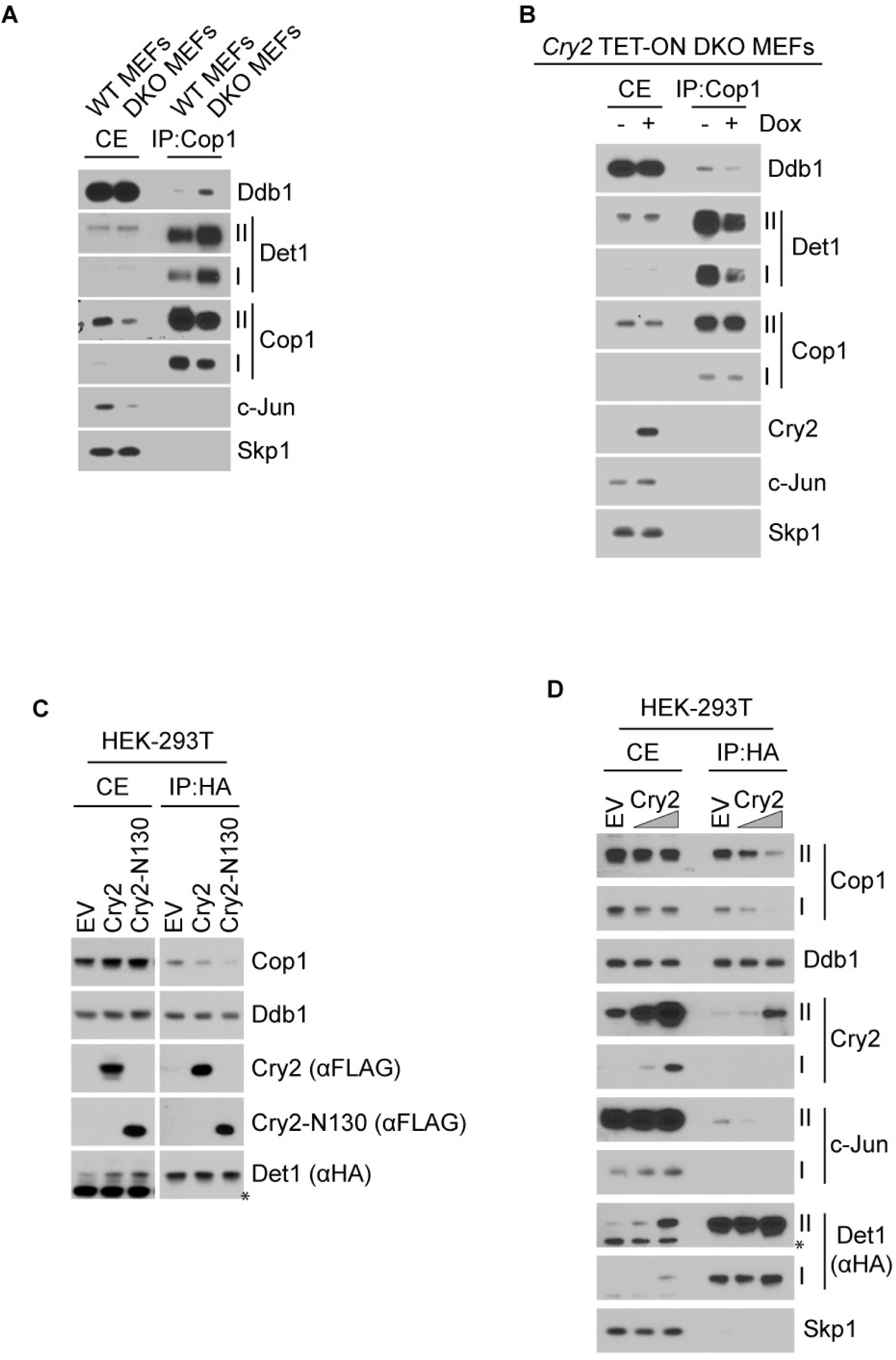
Cry1 and Cry2 inhibit the binding between Cop1 and Det1-Ddb1. **(A)** Endogenous Cop1 immunoprecipitation in wild-type MEFs and DKO MEFs. Cell extracts and immunoprecipitates were analyzed by immunoblotting with antibodies to the indicated proteins. Roman numbers indicate different exposure times: I, short exposure; II, long exposure. **(B)** Endogenous Cop1 immunoprecipitation in DKO MEFs infected with lentiviruses expressing untagged Cry2 under the control of a doxycycline-inducible promoter. Cell extracts and immunoprecipitates were analyzed by immunoblotting with antibodies to the indicated proteins. Human Cry2 was exogenously expressed in DKO MEFs and tested with antibodies against human Cry2. Roman numbers indicate different exposure times: I, short exposure; II, long exposure. **(c)** Exogenously expressed HA tagged Det1 co-expressed with either GFP expressing vector or with Cry2 wild-type expressing vector or a Cry2-N130 truncation mutant expressing vector were transfected in HEK-293T cells followed by immunoprecipitation from cell extracts with anti-HA resin. Cell extracts and immunoprecipitates were analyzed by immunoblotting with antibodies to the indicated proteins. The asterisk indicates a cross-reacting band. **(D)** Exogenously expressed HA tagged Det1 was transfected in HEK-293T cells together with either a GFP expressing vector or increasing amounts of a wild-type Cry2 expressing vector. Next, immunoprecipitation was carried out from cell extracts with an anti-HA resin. Cell extracts and immunoprecipitates were analyzed by immunoblotting with antibodies to the indicated proteins. The asterisk indicates a cross-reacting band. Roman numbers indicate different exposure times: I, short exposure; II, long exposure.

The physical interaction between Cryptochromes and Det1 (Fig. 3A and Fig. 3C), together with the increased interaction between Cop1 and Det1 in absence of the Cryptochromes (Fig. 4A) and the reduced interaction between Cop1 and Det1 after transient expression of either Cry1 or Cry2 (Fig. S4A-B) led us to postulate that Cryptochromes could displace Cop1 from Det1. To test this hypothesis, we immunopurified HA-tagged Det1 in the presence of either a control vector, or wild-type Cry2, wild-type Cry1, Cry2-N130, or Cry1-N112 (the Cry1 truncated mutant equivalent to Cry2-N130; see Fig. S5). Strikingly, co-expression of these constructs, but not the control vector, reduced the binding between Cop1 and Det1 (Fig. 4C and Fig. S4C). Additionally, expression of increasing amounts of exogenous wild-type Cryptochromes resulted in a dose-dependent displacement of endogenous Cop1 from Det1 (Fig. 4D and Fig. S4D). In contrast, expression of Cry2, Cry2-N130, Cry1, and Cry1-N112 did not affect the binding between Det1 and Ddb1 (Fig. 3A, Fig. 4C-D, and Fig. S4C-D).

All together, these results strongly suggest that the interaction between Cryptochromes and Det1 displaces Cop1 from Det1 and the rest of the CRL4 complex, which could explain the effect of Cryptochromes on the stabilization of Cop1 substrates.

Contrary to the current assumption, our results indicate that the Cryptochromes-Cop1 axis is present in mammals. The similarity between the axes in plants and mammals comes from the Cryptochromes-mediated inactivation of the CRL4^Cop1^complex and the consequent stabilization of Cop1’s substrates. However, the molecular mechanisms of this inactivation, as well as the signaling events downstream of the Cryptochromes-Cop1 axis differ from plants to mammals. Whereas in plants Cryptochromes bind constitutively to Cop1 and inhibit its activity in a blue light-dependent manner (leading to the accumulation of its substrates), we found that in mammals Cryptochromes bind Det1, instead of Cop1, and block the ability of Cop1 to interact with the CRL4 complex, leading to the accumulation of CRL4^Cop1^ substrates (see model in Fig. S4E).

We tested the Cryptochromes-Cop1 axis in the context of the well-established Cryptochromes-mediated repression of the GR transcriptional network, finding that, both in MEFs and *in vivo,* Cryptochromes repress GR, at least in part, through the inactivation of Cop1. Among the numerous CRL4^Cop1^ substrates, we focused on c-Jun, which is both a canonical and constitutive substrate of Cop1, as well as a repressor of the GR transcriptional output [3, 20–23]. We found that Cryptochromes positively control c-Jun protein levels by inhibiting Cop1. Moreover, our motif enrichment analysis shows an overrepresentation of c-Jun and Fos recognition motifs at the promoter of genes regulated by dexamethasone in combination with either Cryptochromes’ stabilization or Cop1 silencing. Although Cryptochromes repress GR-dependent transcription, it has remained puzzling the fact that they have no effect on the GR-dependent repression of the NF-⍰ B inflammatory gene network [9]. We propose that Cryptochromes repress GR via the Cop1-c-Jun axis, explaining why Cryptochromes do not interfere with the NF-⍰ B-dependent transcription.

We studied the effect of the Cryptochromes-Cop1-GR axis on the GR transcriptional activity in cell systems and mouse liver. However, Cop1 has an ever-growing number of substrates, raising the possibility that the Cryptochromes-Cop1 axis controls a vast signaling network. Thus, our study opens the door to future investigations on the Cryptochromes-dependent stabilization of Cop1 substrates and their specific roles in different organs and tissues.

## Materials and Methods

### Cell Culture

WT immortalized MEFs (iMEFs) and CRY1^-/-^;CRY2^-/-^ iMEFs (DKO MEFs) from littermate mice were kindly provided by Dr. Choogon Lee. c-Jun^-/-^ iMEFs were kindly provided by Dr. Michael Karin. Beta TC-6 cells were kindly provided by Dr. Joseph Bass. HEK293T cells and U-2OS cells were obtained by the American Type Culture Collection. CRY1^-/-^;CRY2^-/-^ iMEFs and Beta TC-6 cells stably infected with pTRIPZ vectors were propagated in DMEM supplemented with 10% Tet system-approved FBS (Takara/Clontech Laboratories). Doxycycline (Sigma-Aldrich) was used at 100 ng/mL. All the other cell lines were maintained in Dulbecco’s modified Eagle’s medium containing 10% fetal bovine serum (Corning Life Sciences). MG132 (Peptides International) at 10 µM. Cells were periodically screened for *Mycoplasma* contamination.

### Plasmids

CRY1 was a gift from Aziz Sancar (Addgene plasmid # 25843), CRY2 was a gift from Aziz Sancar (Addgene plasmid # 25842), DET1 pcDNA4/TO/myc-His (Abgent cat. No. DC09517). cDNAs were subcloned in pcDNA 3.1 (Life Technologies) and in pTRIPZ (GE healthcare). c-Jun cDNA was amplified by PCR using a cDNA library generated from HEK293T cells and sub-cloned into pBabe (Cell Biolabs). Human CRY2 mutants were generated by using the QuikChange Site-directed Mutagenesis kit (Stratagene). All cDNAs were sequenced.

### Gene Silencing by siRNA

ON-TARGETplus SMARTpool siRNA oligos targeting either COP1 or TRIP6 were transfected using RNAi Max (Dharmacon). ON-TARGETplus non-targeting siRNA (Dharmacon, catalog no. D-001810-01) served as a negative control.

### Retro- and Lentivirus-Mediated Gene Transfer

HEK-293T cells were transiently co-transfected with retroviral (pBabe) vectors containing vesicular stomatitis virus G protein (VSV-G) and the gene of interest along with pCMV-Gag-Pol using polyethylenimine. Alternatively, lentivirus (pTRIPZ) vectors containing vesicular stomatitis virus G protein (VSV-G) and the gene of interest along with pCMV Delta R8.2 were co-transfected using polyethylenimine. Retrovirus- or lentivirus-containing medium, 48 hr after transfection, was collected and supplemented with 8 mg/ml Polybrene (Sigma). Cells of interest were then infected by replacing the cell culture medium with the viral supernatant for 6 hours. Selection of stable clones was carried out with puromycin.

### Immunoprecipitation and immunoblotting

HEK-293T cells were transiently transfected using polyethylenimine (PEI). U-2OS cells were transiently transfected using siLentFect (Bio-rad). Cell lysis was carried out with lysis buffer (50 mM Tris pH 8.0, 150 mM NaCl, 10% glycerol, 1 mM EDTA, and 0.5% NP-40) supplemented with protease and phosphatase inhibitors. Immunoprecipitations were carried out after 3 hours treatment with MG132 and with either an anti-FLAG antibody conjugated to agarose resin or an anti-HA antibody conjugated to agarose resin. For immunoprecipitation of endogenous proteins, wild-type iMEFs were collected and lysed with lysis buffer. Cop1 was immunoprecipitated with the listed antibody mixed with Protein A sepharose beads (Invitrogen). Elution of the immunoprecipitate was carried out with NuPAGE® LDS sample buffer (Thermo Fisher Scientific) supplemented with β-mercaptoethanol (Sigma-Aldrich) and incubation at 95°C for 3 minutes. The following antibodies were used: Cop1 (1:2000, Bethyl #A300-894A), Cry1 (1:2000, Bethyl #A302-614A), c-Jun (1:2000, Cell signaling #9165s), Skp1 (1:5000, produced in our laboratory), for detection of human CRY2 (1:2000, Bethyl #A302-615A), for detection of mouse CRY2 (1:2000, Proteintech #13997-1-AP). JunB (1:2000, Bethyl #A302-704A), Actb (1:7000, Sigma-Aldrich #A5441), Ets-1 (1:2000, Cell signaling #14069), p53 (1:1000, Santa Cruz Biotechnology #sc-6243), FLAG (1:7000, Sigma-Aldrich #F7425), Stat3 (1:2000, Santa Cruz Biotechnology #sc-8019), Mta1 (1:2000, Santa Cruz Biotechnology #sc-17773), Atgl (1:2000, Santa Cruz Biotechnology #sc-365278), Ddb1 (1:5000, Bethyl #A300-462A), HA (1:2000, Bethyl #A190-108A), Per2 (1:2000, Bethyl #A303-109A), Clock (1:2000, Bethyl #A302-618A), Bmal1 (1:2000, Bethyl #A302-616A), Fbxl3 (1:2000, Signalway antibody #21459), Det1 (1:500, Santa Cruz Biotechnology #sc-514348), Cul4 (1:2000, Bethyl #A300-793A), Dda1 (1:2000, Proteintech #14995-1-AP).

### qRT-PCR

Total RNA was generated using RNeasy mini kits (Qiagen). cDNA was generated using Random Hexamers EcoDry kits (Takara Clontech). qPCR was performed using Absolute SYBR green (Thermo Fisher Scientific) on a Roche Lightcycler 480. Analysis of the qPCR experiments was conducted via absolute relative quantification with in-experiment standard curves for each primer set to control for primer efficiency. The oligos used for qRT-PCR analysis are listed:

Quantitative PCR primer sequences for cell cultures experiments

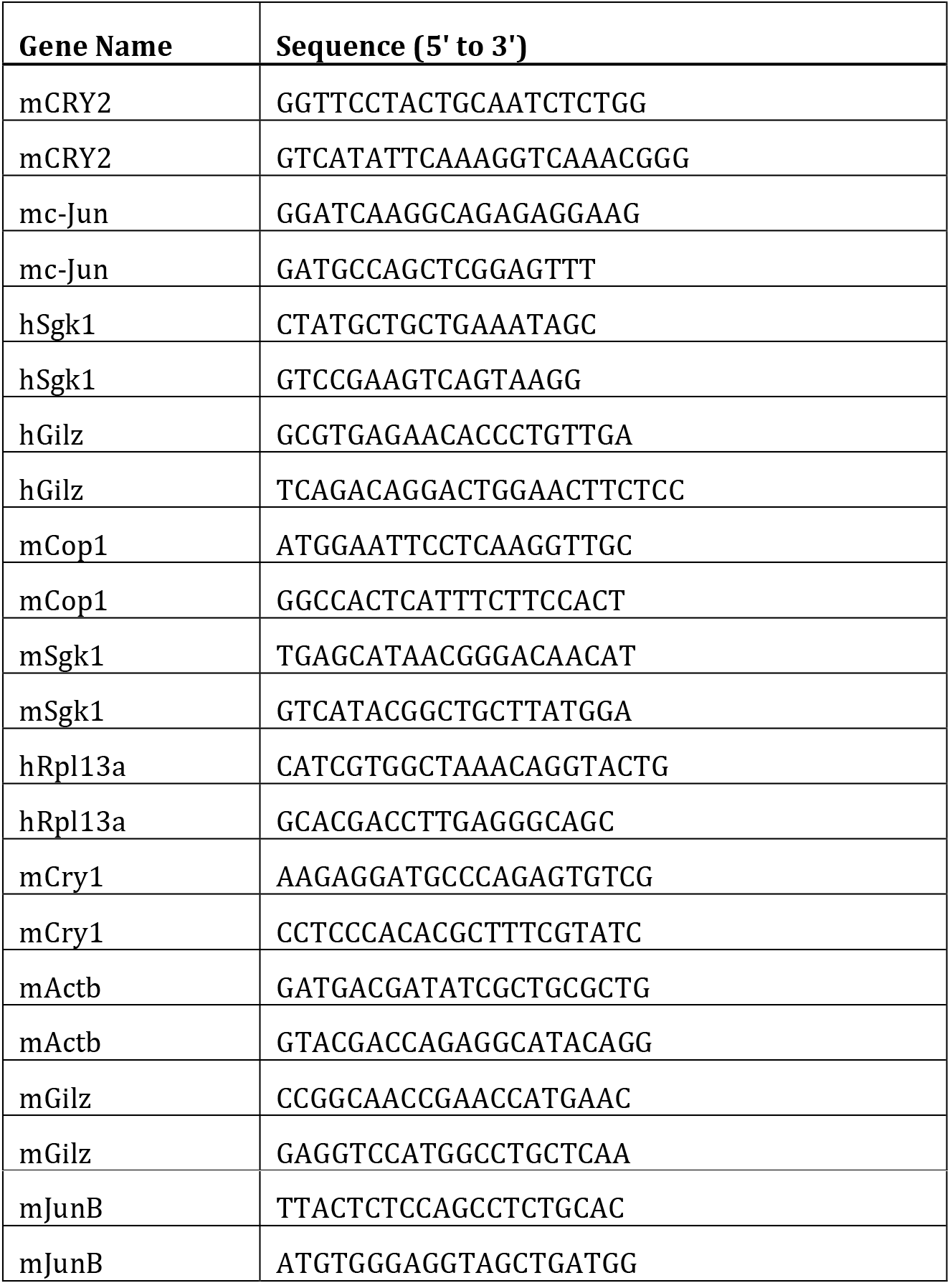

### Mass spectrometry

HEK-293T cells were transfected with constructs encoding either FLAG-tagged human COP1, co-transfected with GFP control vector, or human wild-type CRY1, or human wild-type CRY2. Forty-eight hours after transfection, cells were collected and lysed in lysis buffer (50 mM Tris-HCl pH 7.5, 150 mM NaCl, 1 mM EDTA, 0.5% NP-40, plus protease and phosphatase inhibitors). Proteins were immunopurified with anti-FLAG M2 agarose beads (Sigma) and, after extensive washing, eluted by competition with 3x-FLAG peptide (Sigma). The eluate was then precipitated with TCA. Next, mass-spectrometric analysis of the immunoprecipitates was carried out according to (*32*).

### RNA Extraction and Next-Generation Sequencing Analysis

Libraries for RNA-seq were prepared according to manufacturer’s instructions (Illumina). Briefly, total RNA was extracted from cultured cells using the RNeasy Plus Mini Kit (QIAGEN). Poly(A) RNA was isolated using Dynabeads Oligo(dT)25 (Invitrogen) and used as input for library construction utilizing the dUTP method. Barcodes were used for sample multiplexing. RNA libraries were sequenced on an Illumina HiSeq. RNAseq differential expression analysis was performed for three lanes of a single-read 50 Illumina HiSeq 2500 run. Per-sample FASTQ files were generated using the bcl2fastq Conversion software (v.1.8.4) to convert per-cycle BCL base call files outputted by the sequencing instrument into the FASTQ format. The alignment program, STAR (v2.4.5a), was used for mapping reads of 24 mouse samples to the mouse reference genome mm10 and the application FastQ Screen (v0.5.2) was utilized to check for contaminants. The software, featureCounts (Subread package v1.4.6-p3), was used to generate the matrix of read counts for annotated genomic features. For the differential gene statistical comparisons between groups of samples contrasted by non-targeting RNA interference and targeting Cop1 RNA interference conditions as well as KL001 and dexamethasone chemical treatments, the DESeq2 package (Bioconductor v3.3.0) in the R statistical programming environment was utilized. The heatmap was generated using the function ’heatmap.2’ within the package ’gplots’ in the R statistical programming environment.

### Transcription factor motif analysis

Motifs enriched within promoters of dexamethasone-inducible genes whose expression was modified by treatment with KL001 and COP1 siRNA were identified using the Homer script “findMotifs.pl” (PMID 20513432). Specifically, we supplied the gene IDs for either the 396 dexamethasone-regulated genes that were also differentially regulated by KL001 and siCOP1 treatment or the remaining 3144 dexamethasone-sensitive genes and searched for motifs among these two groups within −300 to +50 bp of the annotated transcription start site and allowing for at most two mismatched bases to known or de novo motif assignments.

### Animals

All animal care and use procedures were in accordance with guidelines of the Institutional Animal Care and Use Committee. All experiments were performed using male C57BL/6J mice between 3-4 months of age, and mice were maintained on a 12:12 light dark (LD) cycle in the Northwestern University Center for Comparative Medicine.

### *In vivo* delivery of shRNA via adeno-associated virus (AAV)

AAV-technology was utilized to delivery shRNAs to liver tissues *in vivo*. AAVs were cloned, packaged into serotype 8, purified by Cesium Chloride centrifugation, concentrated to ~1×10^12^ GC/mL, and buffer exchanged to PBS w/5% glycerol by Vector Biolabs (Malvern, PA).Pre-validated shRNAs targeting mRFWD2 (Cop1, NM_011931) or a scrambled control were expressed under the U6 promoter and GFP was co-expressed under the CMV promoter. Mice were anaesthetized using isoflurane inhalation and 1×10^11^ GC (100 µL) of AAV expressing either shCop1 or scrambled control RNAs were delivered by retroorbital injection. After 4 weeks, mice were injected with 100 ⍰ l saline alone or 1 ⍰g water-soluble dexamethasone (Sigma) resuspended in saline one hour prior to being sacrificed at ZT2 and ZT10. Liver tissue was excised and snap frozen in liquid nitrogen. Total RNA was extracted from frozen liver tissue using Tri-Reagent (Molecular Research Center, Inc), and quantitative PCR was performed and analyzed using a CFX384 (Bio-Rad). PCR conditions were: 10 min at 95°C, then 35 cycles of 10 s at 95°C, 15 s at 60°C. Relative expression levels (normalized to *β-actin*) were determined using the comparative CT method.

Quantitative PCR primer sequences for mouse liver tissue experiments

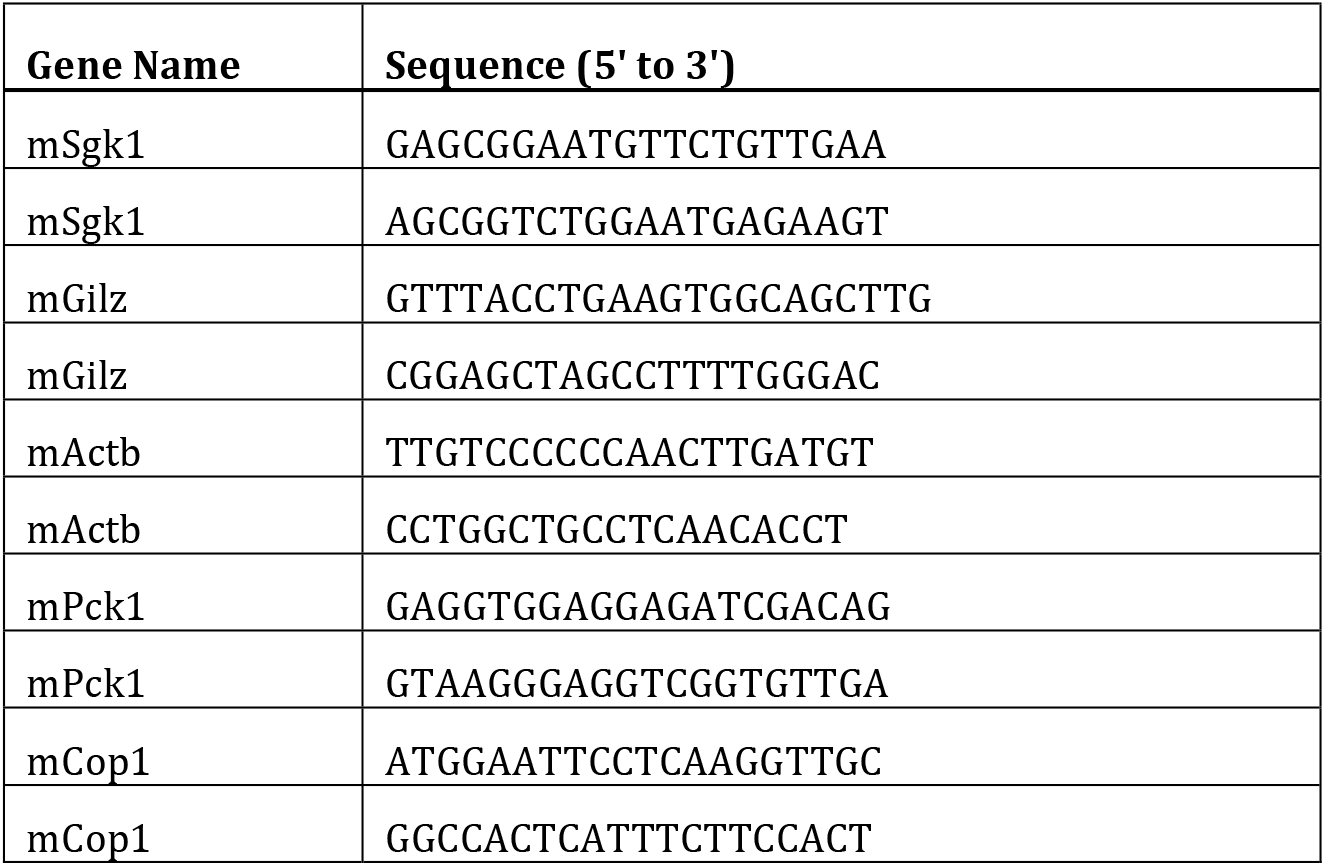

### Statistics and reproducibility

All data were collected and analyzed by Prism 5 (Graphpad). Sample sizes and reproducibility for each figure are denoted in the figure legends. Unless otherwise noted, data are representative of at least three biologically independent experiments. Two-group datasets were analyzed by student’s unpaired T-test. For three- or four-group analysis, one-way ANOVA was used. All graphs show mean values. Error bars indicate +/- S.D. or S.E.M., as indicated in figure legends.

## Data availability

The next-generation sequencing data that support the findings of this study in Figure 1 and in Supplementary figure 1 have been deposited in the Gene Expression Omnibus (GEO) database under the accession code GSE124388. All other data supporting the findings of this study are available from the corresponding author on reasonable request.

**Fig. S1.**
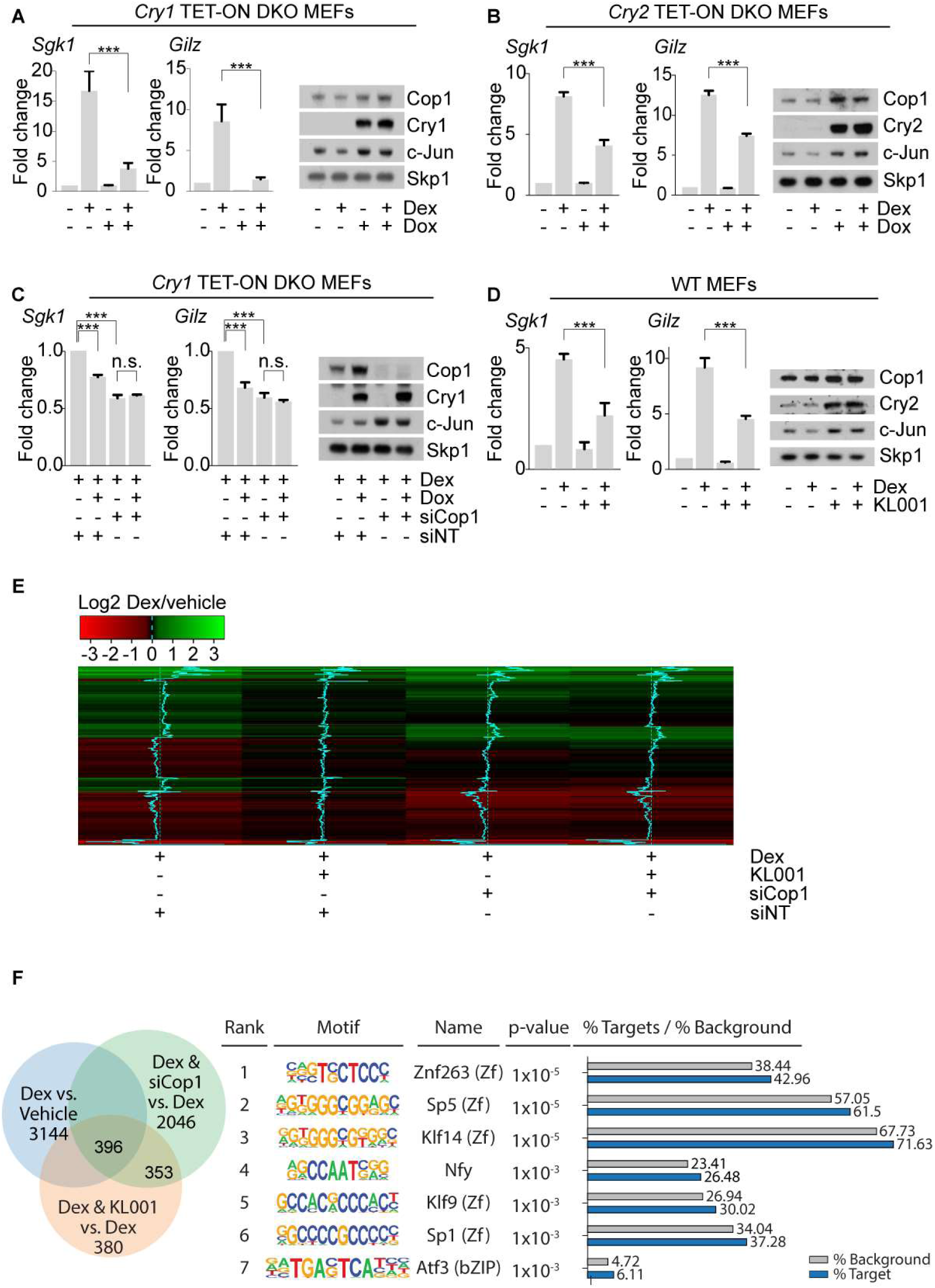
Cop1 mediates the Cryptochromes-dependent rhythmic repression of the GR. **(A)** On the left, qPCR analysis of cDNA prepared from DKO MEFs infected with lentiviruses expressing untagged Cry1 under the control of a doxycycline-inducible promoter and before and after dexamethasone treatment. On the right, cells extracts were analyzed by immunoblotting with antibodies to the indicated proteins. Human Cry1 was exogenously expressed in DKO MEFs. **(B)** On the left, qPCR analysis of cDNA prepared from DKO MEFs infected with lentiviruses expressing untagged Cry2 under the control of a doxycycline-inducible promoter and before and after dexamethasone treatment. On the right, cells extracts were analyzed by immunoblotting with antibodies to the indicated proteins. Cry2 was exogenously expressed in DKO MEFs and tested with antibodies against human Cry2. **(C)** On the left, qPCR analysis of cDNA prepared from DKO MEFs infected with lentiviruses expressing untagged Cry1 under the control of a doxycycline-inducible promoter, after dexamethasone treatment, in presence or absence of Cop1 knockdown. On the right, cells extracts were analyzed by immunoblotting with antibodies to the indicated proteins. Human Cry1 was exogenously expressed in DKO MEFs and tested with antibodies against human Cry1. **(D)** On the left, qPCR analysis of cDNA prepared from wild-type MEFs, before and after KL001 treatment and in presence or absence of dexamethasone treatment. On the right, cells extracts were analyzed by immunoblotting with antibodies to the indicated proteins. **(E)** Dexamethasone-mediated activation of the GR in wild-type MEFs treated with KL001, in presence or absence of Cop1 knockdown. RNA was isolated from the samples and quantified by RNA-seq. Differential expression analysis of RNA-seq data sets, [see next-generation sequencing data deposited in Gene Expression Omnibus (GEO) database under the accession code GSE124388] is represented in the heatmap as log2 fold change for 792 genes. Dexamethasone-deregulated genes were considered differentially expressed when the p-adjusted value was <0.05. (n=3 biologically independent experiments). **(F)** In the left panel, Venn diagram representing differentially regulated genes after dexamethasone treatment alone or in combination with either siRNA-mediated depletion of Cop1 or KL001 treatment in wild-type MEFs. RNA was isolated from the samples and quantified by RNA-seq. [see next-generation sequencing data deposited in Gene Expression Omnibus (GEO) database under the accession code GSE124388]. Regulated genes were considered differentially expressed when the p-adjusted value was <0.05. In the right panel, the table contains the first 7 most highly overrepresented motifs within the promoter region of the 3144 deregulated genes by dexamethasone after subtraction of the 396 common regulated genes among all the tested conditions.

**Fig. S2.**
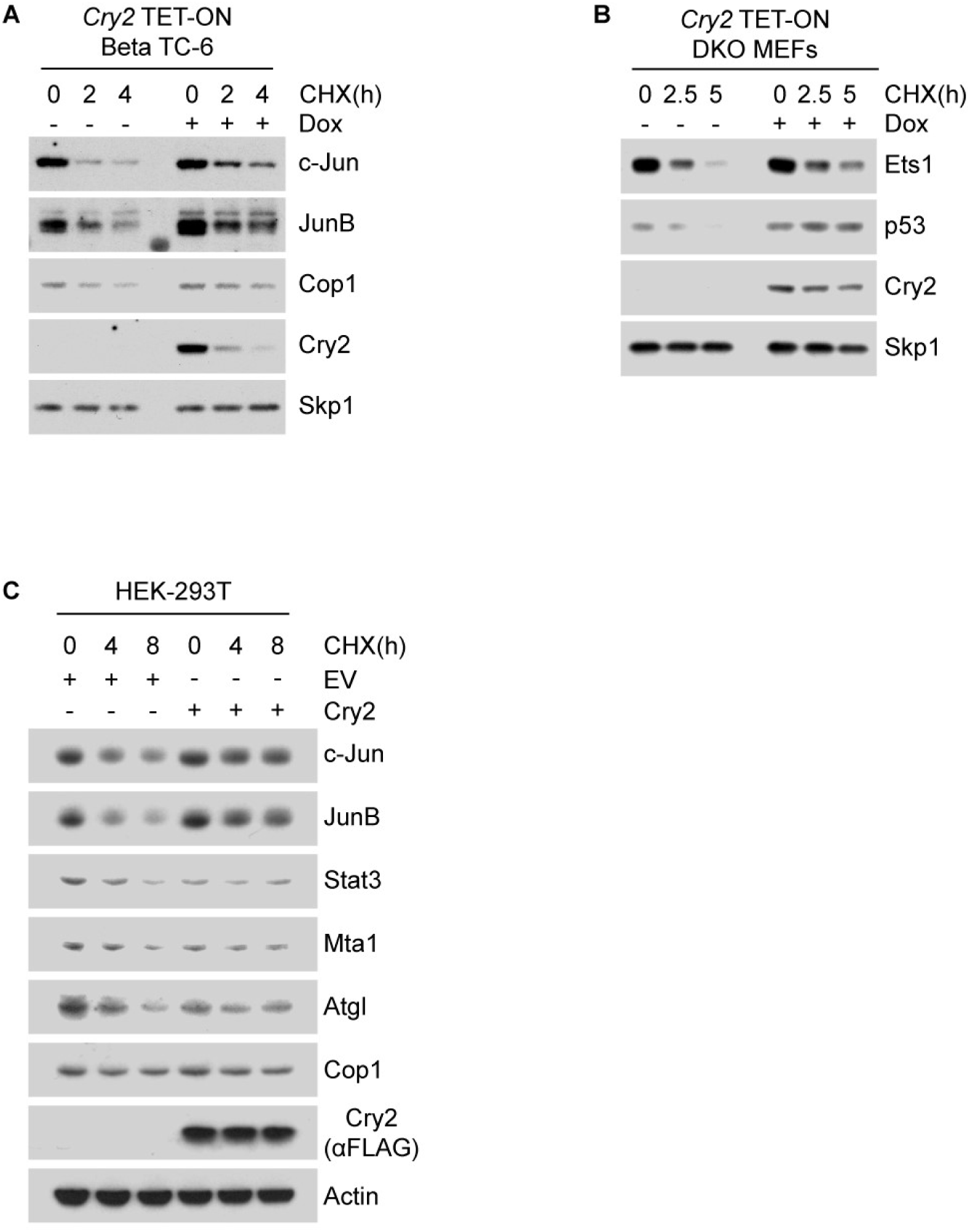
Cry2 overexpression increases the half-life of Cop1 substrates in various cell lines. **(A)** Beta TC-6 cells were infected with lentiviruses expressing untagged Cry2 under the control of a doxycycline-inducible promoter and treated overnight with doxycycline. The next day cycloheximide was applied to the cells for the indicated time and cells were then collected, lysed and immunoblotted with antibodies to the indicated proteins. Human Cry2 was exogenously expressed in Beta-TC-6 cells and tested with the antibody against human Cry2. **(B)** The Western Blot represents the experiment shown in Fig. 2C immunoblotted also with antibodies to Ets1 and p53. Cry2 and Skp1 are shown again as references. **(C)** HEK-293T cells were transfected with either a control GFP expressing vector or a Cry2 wild-type expressing vector. The next day cycloheximide was applied to the cells for the indicated time and cells were then collected, lysed and immunoblotted with antibodies to the indicated proteins.

**Fig. S3.**
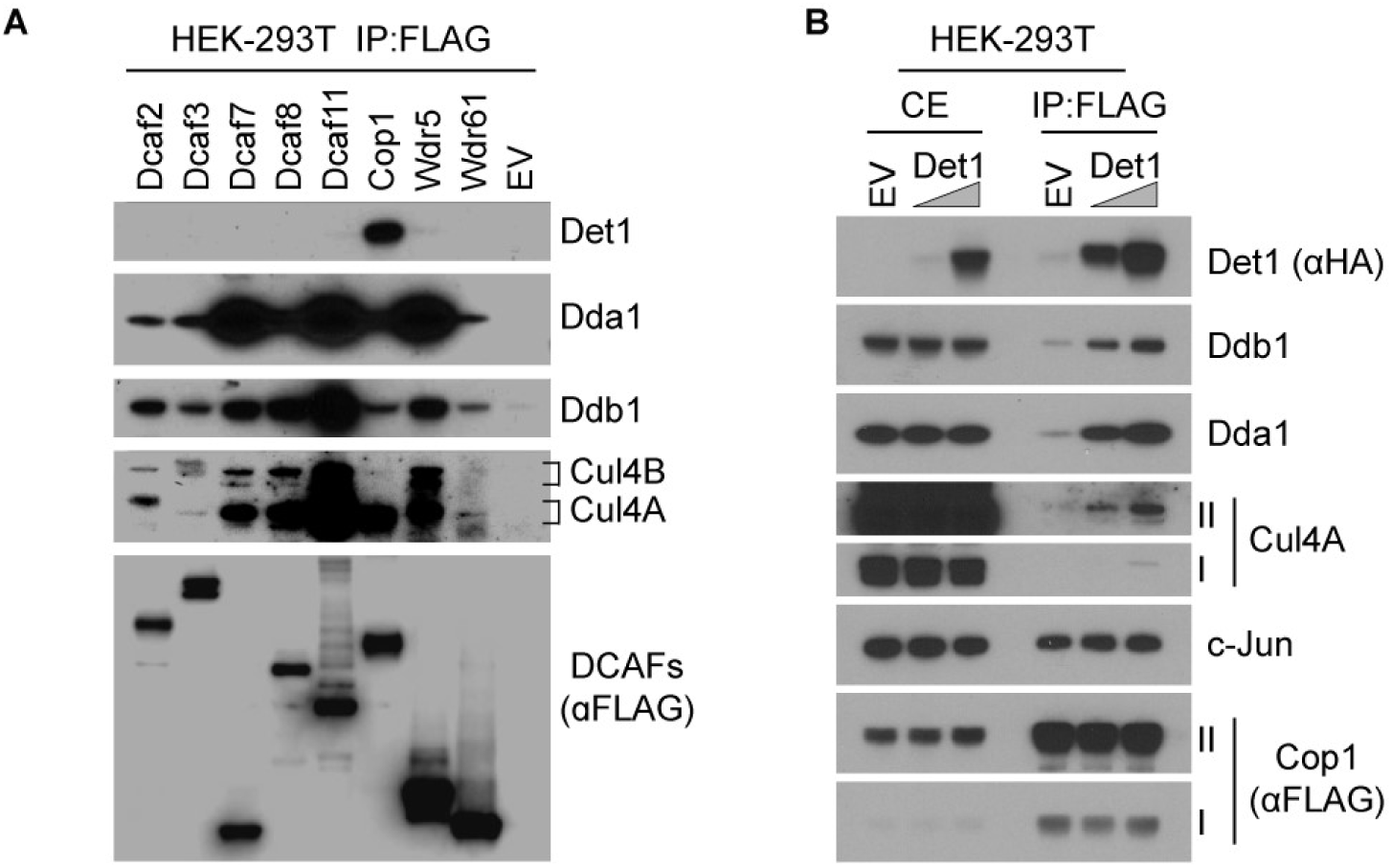
Det1 binds specifically to Cop1 and is rate-limiting for the formation of the CRL4^Cop1^ complex. **(A)** HEK-293T cells were transfected with eight different FLAG tagged DCAFs followed by immunoprecipitation with anti-FLAG resin and analysis by immunoblotting with antibodies to the indicated proteins. **(B)** HEK-293T cells were transfected with FLAG tagged Cop1 together with increasing doses of HA tagged Det1 followed by immunoprecipitation with anti-FLAG resin and analysis by immunoblotting with antibodies to the indicated proteins. Roman numbers indicate different exposure times: I, short exposure; II, long exposure.

**Fig. S4.**
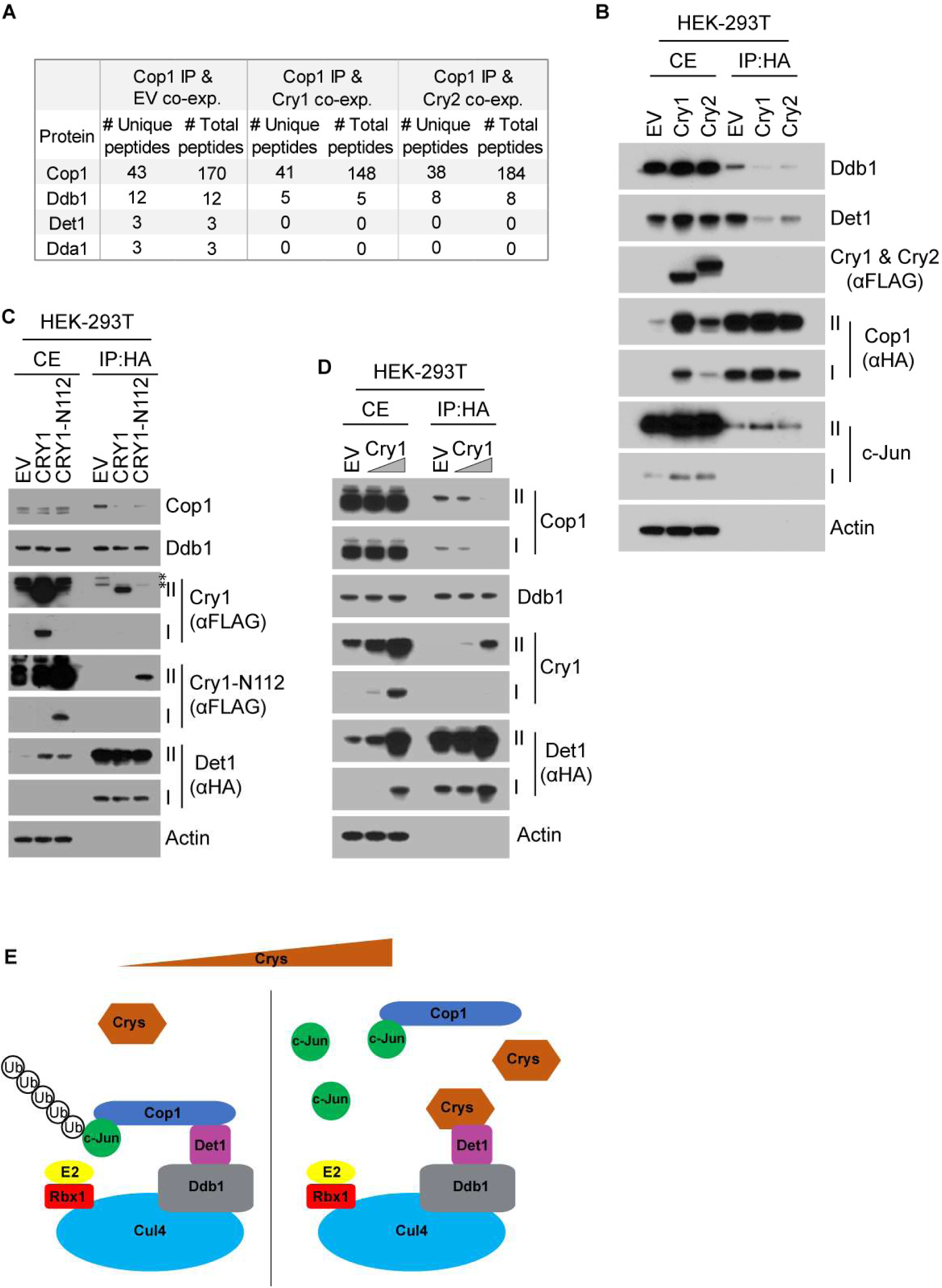
Cry1 and Cry2 inhibit the binding between Cop1 and Det1-Ddb1. **(A)** Number of peptides detected after mass-spectrometric analysis of proteins present in the Cop1 complex, after expression and immunoprecipitation of FLAG tagged Cop1 with anti-FLAG resin in HEK-293T cells. Cop1 was co-expressed with either a GFP control vector or with Cry1 or Cry2 expressing vectors. **(B)** HEK-293T cells were transfected with FLAG tagged Cop1 together with a GFP control vector or with either Cry1 or Cry2 expressing vectors followed by immunoprecipitation with anti-FLAG resin and analysis by immunoblotting with antibodies to the indicated proteins. Roman numbers indicate different exposure times: I, short exposure; II, long exposure. **(C)** Exogenously expressed HA tagged Det1 co-expressed with either GFP expressing vector or with Cry1 wild-type expressing vector or a Cry1-N112 truncation mutant expressing vector were transfected in HEK-293T cells followed by immunoprecipitation from cell extracts with anti-HA resin. Cell extracts and immunoprecipitates were analyzed by immunoblotting with antibodies to the indicated proteins. The asterisks indicate cross-reacting bands. Roman numbers indicate different exposure times: I, short exposure; II, long exposure. **(D)** Exogenously expressed HA tagged Det1 was transfected in HEK-293T cells together with either a GFP expressing vector or increasing amounts of Cry1 wild-type. Next, immunoprecipitation was carried out from cell extracts with an anti-HA resin. Cell extracts and immunoprecipitates were analyzed by immunoblotting with antibodies to the indicated proteins. Roman numbers indicate different exposure times: I, short exposure; II, long exposure. **(E)** Working model. The oscillating levels of the Cryptochromes control the stabilization of Cop1’s substrates by inhibiting the formation of the CRL4^Cop1^ complex.

**Fig. S5.**
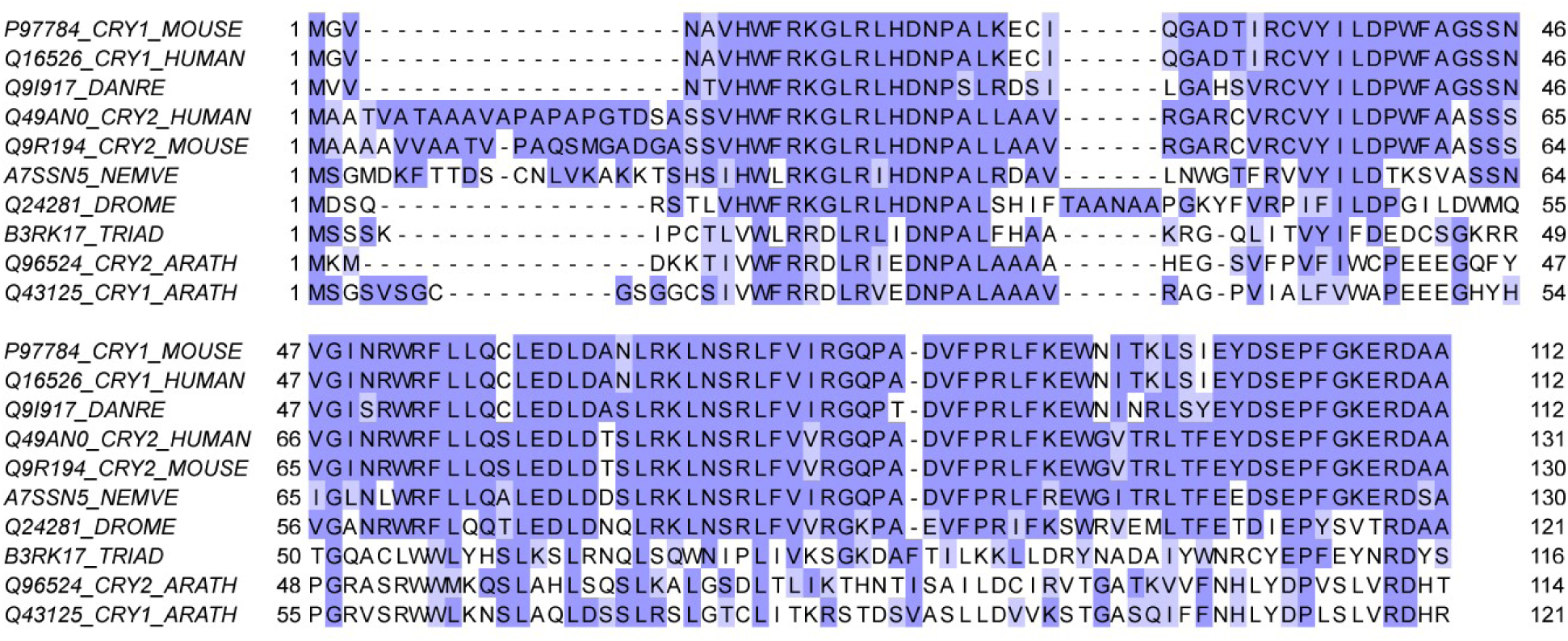
Alignment of the N-termini of Cryptochrome proteins. The N-terminal regions of mammalian Cry1 and Cry2 are well conserved. UniProt Knowledgebase (UniProtKB) identifiers are given for each protein used in the alignment. Representative genomes were chosen: HUMAN (*Homo sapiens*), MOUSE (*Mus musculus*), DANRE (*Danio rerio*), NEMVE (*Nematostella vectensis*), DROME (*Drosophila melanogaster*), TRIAD (*Trichoplax adhaerens*), ARATH (*Arabidopsis thaliana*). The multiple sequence alignment was generated by Tcoffee as an option of the multiple alignment editor Jalview version 2.10.5 (http://www.jalview.org/).

